# GABAergic projection from the subiculum to the medial entorhinal cortex in mice and rats

**DOI:** 10.64898/2026.06.06.730551

**Authors:** Takahiro Aimi, Takumi Shibuya, Haruka Umeno, Keiko Karasawa, Ken-Ichiro Tsutsui, Shinya Ohara, Takuma Kitanishi

## Abstract

The subiculum (SUB) is a major hippocampal output hub that routes information to cortical and subcortical targets, and its long-range projections are considered excitatory. Using enhancer-driven adeno-associated viral vectors to selectively label γ-aminobutyric acid (GABA)-releasing neurons across species, here we show that the dorsal SUB also sends an inhibitory projection to the dorsal part of the medial entorhinal cortex (MEC) in mice and rats. Anterograde tracing in mice revealed that the dorsal SUB contains GABAergic neurons that project sparsely to all layers of the dorsal MEC with enrichment in superficial layers, in contrast to the glutamatergic SUB axons targeting MEC layer V. Slice electrophysiology demonstrated that these GABAergic axons form inhibitory synapses in the MEC. A subset of projecting neurons expressed parvalbumin (PV), whereas somatostatin-positive neurons were rare. Consistently, PV neuron-specific anterograde tracing recapitulated the SUB-to-MEC projection. In rats, subicular GABAergic axons were enriched in MEC layer II, and SynaptoTAG2-labeled presynaptic boutons were positive for the vesicular GABA transporter, supporting inhibitory synapse formation. Anterograde tracing of PV neurons similarly recapitulated the laminar axonal distribution in the MEC. These results identify a conserved PV-associated inhibitory SUB-to-MEC projection with species-specific laminar organization, extending the canonical excitatory view of subicular output.

## Introduction

The subiculum (SUB) is an anatomical and functional hub that transmits hippocampal information to downstream brain regions (Naber and Witter 1998; van Strien et al. 2009; Aggleton and Christiansen 2015; Bohm et al. 2018; Matsumoto et al. 2019). Receiving inputs from the hippocampus and entorhinal cortex, the SUB projects to multiple cortical and subcortical targets, including the entorhinal cortex, retrosplenial cortex, anterior thalamus, nucleus accumbens, and medial mammillary body (Honda and Ishizuka 2015; Winnubst et al. 2019; Umaba et al. 2021). Neurons in the dorsal SUB densely encode multiple navigational variables (Sharp and Green 1994; Lever et al. 2009; Kim et al. 2012; Olson et al. 2017; Gauthier and Tank 2018; Ledergerber et al. 2021; Nakai et al. 2024; Sun et al. 2024) and route this information to downstream targets in a pathway-specific manner (Kitanishi et al. 2021). Individual pathways originating from the dorsal SUB are required for distinct aspects of spatial and contextual memory (Roy et al. 2017; Cembrowski, Phillips, et al. 2018; Yamawaki et al. 2019; Nelson et al. 2020; Yanakieva et al. 2024; Kinman et al. 2025). To date, subicular efferent projections have been reported to arise from excitatory, glutamatergic neurons (Bienkowski et al. 2018; Cembrowski, Phillips, et al. 2018; Cembrowski, Wang, et al. 2018; Ding et al. 2020; Baccini et al. 2025).

The SUB also contains γ-aminobutyric acid (GABA)-releasing neurons, including subsets that express parvalbumin (PV) or somatostatin (SST) (Knopp et al. 2008; Leao et al. 2012; Nichol et al. 2018). However, the properties of the subicular GABAergic neurons remain poorly characterized (Matsumoto et al. 2019). In adjacent hippocampal and parahippocampal regions, diverse GABAergic neuron subtypes have been extensively studied and classified on the basis of distinct features such as gene expression, electrophysiological properties, and/or cellular morphology. Most of these subtypes function as interneurons that form synapses locally within a given brain region and regulate local circuit operations (Freund and Buzsáki 1996; Klausberger and Somogyi 2008; Topolnik and Tamboli 2022; Tzilivaki et al. 2023).

In addition to interneurons, accumulating evidence indicates that specific GABAergic neuron subtypes in various brain regions extend long-range axons beyond their region of origin (Melzer and Monyer 2020). These GABAergic projection neurons can modulate activity in distant brain regions; therefore, they have been proposed to regulate various cognitive functions. For instance, the mouse hippocampal CA1 area and dentate gyrus contain SST-positive GABAergic neurons that project to the medial entorhinal cortex (MEC) (Melzer et al. 2012). Moreover, the MEC sends a backward GABAergic projection to the hippocampus, and this bidirectional connection enhances theta-band rhythmic activity (Melzer et al. 2012). The presubiculum, located adjacent to the SUB, also contains GABAergic neurons that project to the MEC in rats (van Haeften et al. 1997).

Despite the accumulating literature on GABAergic projections, whether the SUB contains such projection neurons has remained unknown. Recently, selective gene expression in GABAergic neurons and their subtypes has been achieved using adeno-associated viral vectors (AAVs) carrying cell type-specific enhancers, which can be applied across vertebrate species (Dimidschstein et al. 2016; Hrvatin et al. 2019; Vormstein-Schneider et al. 2020; Furlanis et al. 2025). By combining enhancer-driven, AAV-based tracing *in vivo* and optogenetic circuit mapping in slice preparations, we demonstrate that the dorsal SUB contains PV-positive GABAergic neurons that project to the dorsal MEC in both mice and rats.

## Results

### Dorsal SUB contains GABAergic neurons projecting to the dorsal MEC in mice

We performed anterograde tracing of GABAergic neurons by injecting AAV expressing green fluorescent protein (GFP) under the control of the GABAergic neuron-selective Dlx enhancer, AAV1-mDlx-GFP (Dimidschstein et al. 2016), into the mouse dorsal SUB (Figs. 1a–e, n = 8 mice). GFP expression was restricted to SUB neurons positive for the 67-kDa isoform of glutamic acid decarboxylase (GAD67), a pan-GABAergic neuronal marker (Fig. 1b). Following localized injection into the dorsal SUB with minimal leakage into adjacent regions (Fig. 1c), we observed many GFP-labeled axons in the dorsal MEC (Figs. 1d and 1e). These GFP-labeled axons were sparsely distributed across all layers of the MEC (Fig. 1e). Quantification indicated that the labeled axons were enriched in the superficial layers. In particular, layers I and III contained more labeled axons than all other layers, whereas layer II had more labeled axons than layer VI (Fig. 1f).

**Fig. 1.**
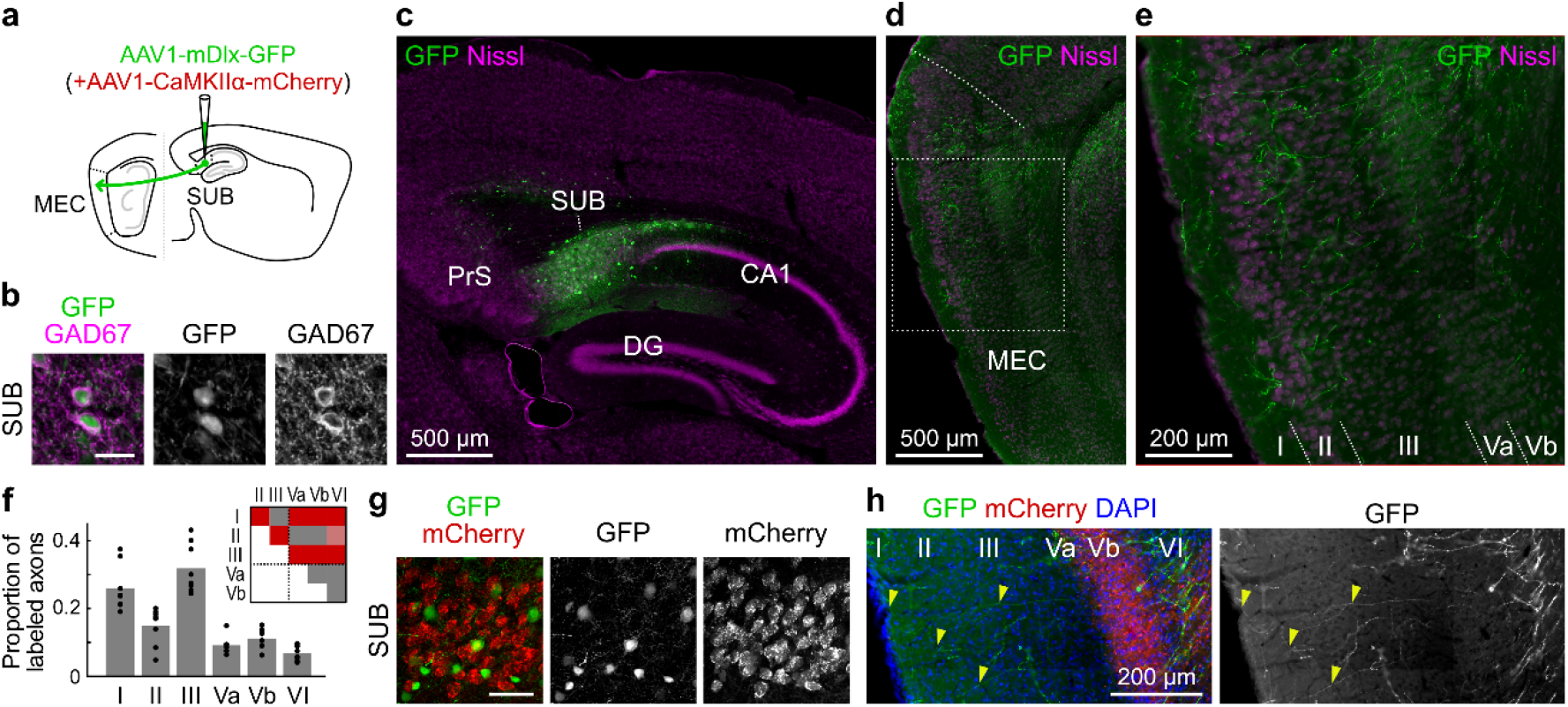
Anterograde tracing of subicular GABAergic neurons in mice. (a) Schematic of the experiment. AAV1-mDlx-GFP alone (b–f) or a mixture with AAV1-CaMKIIα-mCherry (g, h) was injected into the mouse dorsal subiculum (SUB). MEC, medial entorhinal cortex. (b) GFP expression in cells positive for 67-kDa isoform of glutamic acid decarboxylase (GAD67) in the SUB. Scale bar, 20 µm. (b–h) All images are sagittal sections. (c) Representative section near the AAV injection site showing GFP expression in the SUB. PrS, presubiculum; DG, dentate gyrus. (d) GFP-labeled axons in the dorsal part of the MEC ipsilateral to the AAV injection site. Dotted curve, the dorsal boundary of the MEC. (e) High-magnification image of the dorsal MEC (dotted square in d) showing GFP-labeled axons. Numbers, MEC layers; dotted lines, layer boundaries. (f) Proportion of GFP-labeled axon lengths in MEC layers. Dots, individual mice; bars, means. The inset presents a *P*-value table. Red, *P* < 0.01; light red, *P* < 0.05; gray, non-significant; Tukey–Kramer test. Dotted lines, borders between superficial (I–III) and deep (Va–VI) layers. (g) GFP and mCherry expression in the SUB following coinjection. Scale bar, 50 µm. (h) GFP-labeled and mCherry-labeled axons in the dorsal MEC. Yellow arrowheads indicate examples of GFP-labeled axons. Numbers, MEC layers.

SUB glutamatergic neurons innervate the deep layers (especially layer V) of the MEC (Witter et al. 2017; Ohara et al. 2023). To compare the laminar distributions of GABAergic and glutamatergic axons, we coinjected a mixture of AAV1-mDlx-GFP and AAV1-CaMKIIα-mCherry (expressing mCherry under the control of CaMKIIα promoter) into the dorsal SUB (Figs. 1g and 1h, n = 4 mice). GFP and mCherry were expressed in largely nonoverlapping neuronal populations in the SUB (Fig. 1g). Whereas GFP-labeled axons were observed sparsely across all MEC layers, mCherry-labeled axons were predominantly distributed in MEC layer Vb, indicating distinct laminar innervation by subicular GABAergic and glutamatergic neurons.

### Subicular GABAergic neurons preferentially target MEC layer II neurons in mice

To identify the postsynaptic targets of subicular GABAergic inputs in the MEC, we combined optogenetic stimulation of subicular GABAergic axons with whole-cell patch–clamp recordings from MEC neurons. Channelrhodopsin-2 (ChR2)-mCherry was expressed in subicular GABAergic neurons by injecting AAV9-mDlx-ChR2-mCherry into the dorsal SUB. Whole-cell patch–clamp recordings were then performed from MEC neurons in sagittal brain slices while optically stimulating GABAergic axon terminals originating from the dorsal SUB (Fig. 2a, n = 12 mice). Recorded neurons were filled with biocytin and subsequently classified based on the soma location and morphological characteristics. Consistent with previous studies, layer II principal neurons were identified as either stellate cells or pyramidal neurons.

**Fig. 2.**
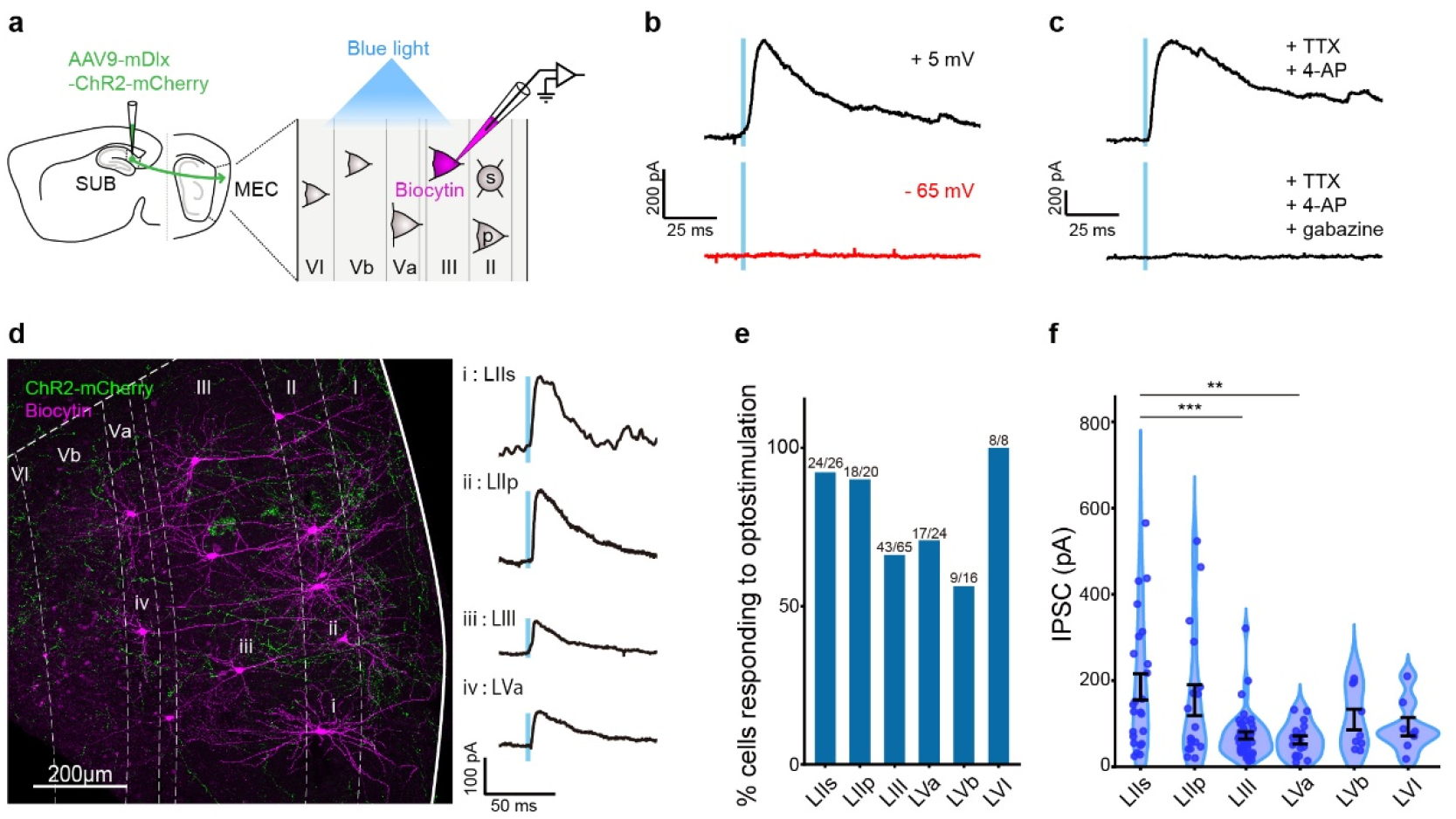
Postsynaptic targets of subicular GABAergic neurons in the mouse MEC. (a) Schematic of the experimental design. AAV9-mDlx-ChR2-mCherry was injected into the dorsal SUB to selectively express ChR2 in subicular GABAergic neurons. Optogenetically evoked synaptic responses were recorded from MEC neurons using whole-cell patch–clamp recordings. (b) PSCs were recorded at holding potentials of +5 mV and −65 mV in response to optical stimulation of subicular GABAergic axon terminals. (c) PSCs recorded at a holding potential of +5 mV in the presence of TTX and 4-AP. Responses were completely abolished by applying gabazine. (d) Representative confocal image of MEC slice showing the distribution of ChR2-mCherry-positive axons (green) and recorded neurons labeled with biocytin (magenta). Optogenetically evoked IPSCs recorded from neurons in the image are presented on the right. (e) Proportion of MEC neurons responding to optical stimulation across layers. (f) IPSC amplitudes in response to optical stimulation across layers (one-way ANOVA, *F*_5,113_ = 5.42, *P* < 0.001; Bonferroni’s multiple comparison test, \*\**P* < 0.01, \*\*\**P* < 0.001.)

We first verified that the optogenetically evoked responses originated from GABAergic axons. Postsynaptic currents (PSCs) were recorded at holding potentials corresponding to the reversal potentials for AMPA receptors (+5 mV) and GABA-A receptors (−65 mV). Robust outward currents were observed at +5 mV, whereas little to no PSCs were detected at −65 mV (Fig. 2b). These optogenetically evoked inhibitory PSCs (IPSCs) persisted in the presence of tetrodotoxin (TTX) and 4-aminopyridine (4-AP), but were abolished by applying the competitive GABA-A receptor antagonist, gabazine (Fig. 2c). Together, these results indicate that the IPSCs recorded at +5 mV reflect monosynaptic GABAergic input originating from the dorsal SUB. We therefore analyzed these IPSCs across different MEC layers.

Anatomical analysis revealed that labeled subicular axons were distributed throughout the dorsal MEC, with a higher density in superficial layers (Fig. 2d). Consistent with this anatomical observation, a high percentage of recorded layer II stellate cells (LIIs, 92%; 24 out of 26 cells) and pyramidal cells (LIIp, 90%; 18 out of 20 cells) responded to optical stimulation (Fig. 2e). In contrast, the proportion of responsive neurons was lower in deeper layers, including layer III (LIII, 66%; 43 out of 65 cells), layer Va (LVa, 71%; 17 out of 24 cells), and layer Vb (LVb, 56%; 9 out of 16 cells). Moreover, even among responsive neurons, IPSC amplitudes in LIII and LVa were significantly smaller than those in layer II stellate cells (Fig. 2f; LII stellate cells, 185.4 ± 30.7 pA; LII pyramidal cells, 154.4 ± 36.1 pA; LIII cells, 72.8 ± 8.4 pA; LVa cells, 62.4 ± 9.3 pA; LVb cells, 109.8 ± 23.9 pA; mean ± SEM). All recorded layer VI neurons (LVI, 8 out of 8 cells) responded to light stimulation. However, consistent with the relatively sparse axonal distribution, IPSC amplitudes were comparable to those in other deep-layer neurons (92.9 ± 21.3 pA). These results demonstrate that subicular GABAergic neurons preferentially provide strong monosynaptic inhibitory input to MEC layer II neurons, particularly stellate cells.

### Dorsal SUB contains PV neurons projecting to the dorsal MEC in mice

Because both the hippocampal CA1 area and the presubiculum, the adjacent areas of the SUB, contain GABAergic projection neurons that innervate the MEC (van Haeften et al. 1997; Melzer et al. 2012), we sought to verify that the GABAergic projection to the MEC originates from the SUB. To this end, we used a dual AAV approach to selectively label MEC-projecting subicular GABAergic neurons, enabling the identification of their somatic locations and molecular subtypes. The retrogradely infecting AAV expressing Cre recombinase, AAV6-pgk-Cre (Kitanishi and Matsuo 2017), and the Cre-dependent, GABAergic neuron-selective reporter AAV, AAV1-hDlx-Flex-GFP (Dimidschstein et al. 2016), were injected into the dorsal MEC and dorsal SUB, respectively (Fig. 3a, n = 9 mice). GAD67-positive, GFP-labeled somata were observed within the SUB cell layer (Fig. 3b). Such neurons were not detected in a negative control in which AAV6-pgk-Cre injection was omitted (n = 4 mice). Among the GFP-labeled cells in the SUB cell layer, 24.0% were PV-positive (Fig. 3c), and 4.1% were SST-positive; the subtype was unidentified for the rest of the cells. The GAD67-positive, GFP-labeled somata were distributed throughout the SUB cell layer, with a higher density toward the distal part (Fig. 3d).

**Fig. 3.**
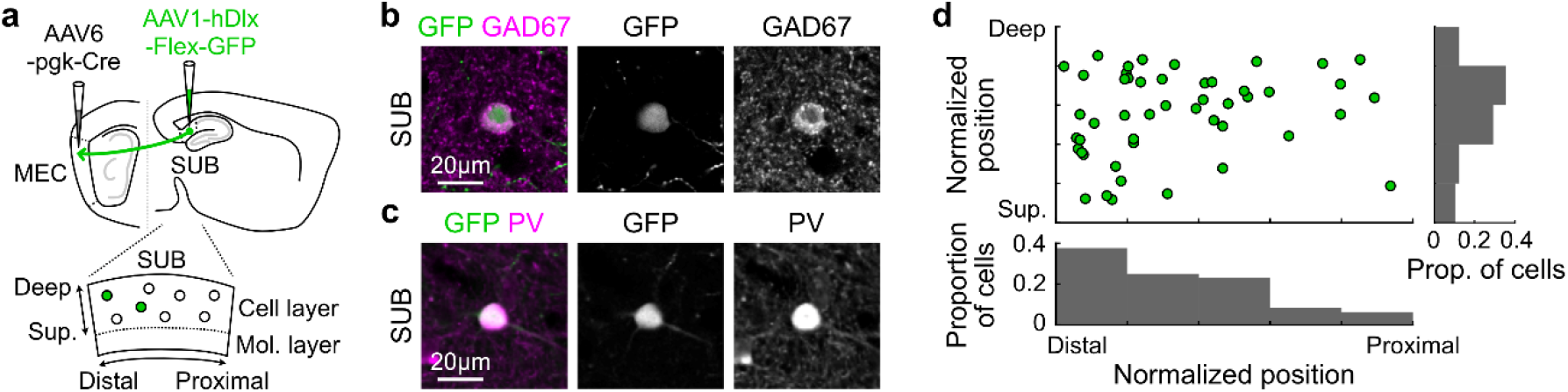
Subtype and distribution of MEC-projecting subicular GABAergic neurons in mice. (a) Schematic of the experiment. AAV6-pgk-Cre and AAV1-hDlx-Flex-GFP were injected into the dorsal MEC and dorsal SUB, respectively, to label the somata of MEC-projecting SUB GABAergic neurons. (b, c) Expression of GAD67 (b) and parvalbumin (PV) (c) in GFP-positive SUB neurons. (d) Distribution of GFP/GAD67 double-positive neurons in the SUB cell layer parallel to the sagittal plane. Distal, away from the CA1 area; proximal, near the CA1 area. Deep, near the alveus; superficial, near the hippocampal fissure. Proportions of cells were nonuniform in both axes: distal-proximal, *χ*^2^ (4) = 15.96, *P* = 0.003; superficial-deep, *χ*^2^ (4) = 12.62, *P* = 0.013; chi-square goodness-of-fit test.

We next performed anterograde tracing of the subicular PV neurons by injecting AAV1-S5E2-GFP-fGFP, which drives GFP expression via a PV neuron-selective E2 enhancer (Vormstein-Schneider et al. 2020), into the dorsal SUB (Fig. 4a, n = 9 mice). Among the GFP-labeled SUB neurons, 84.0% were PV-positive (Fig. 4b). After a localized injection in the SUB with minimal leakage into adjacent regions, we observed GFP-labeled axons sparsely distributed across all dorsal MEC layers (Fig. 4c), similar to the distribution obtained with AAV1-mDlx-GFP (Fig. 1). These results indicate that the dorsal SUB contains PV-positive GABAergic neurons that project to the dorsal MEC in mice.

**Fig. 4.**
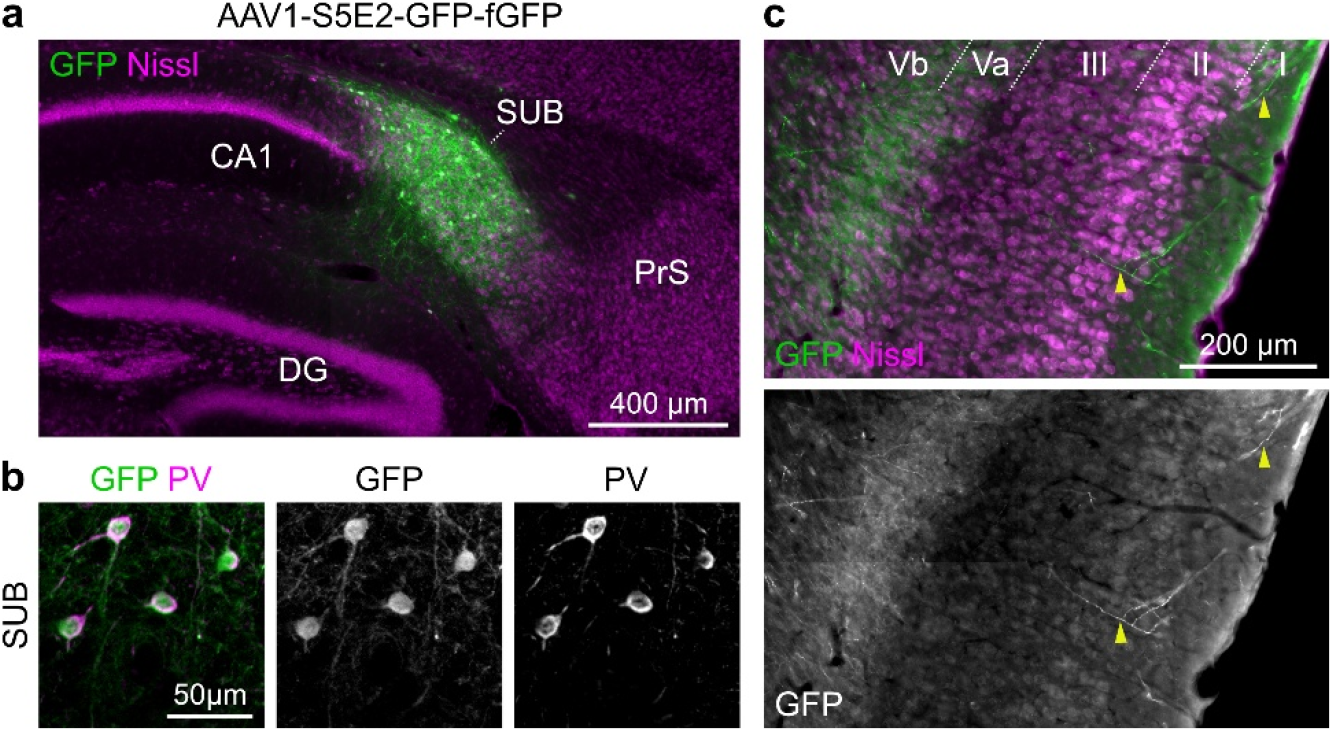
Anterograde tracing of subicular PV neurons in mice. (a) Sagittal section around the dorsal SUB near the AAV1-S5E2-GFP-fGFP injection site. (b) PV expression in GFP-labeled SUB neurons. (c) Sagittal section presenting GFP-labeled axons in the dorsal MEC ipsilateral to the injection site. Numbers, MEC layers; dotted lines, layer boundaries.

### GABAergic projection from the dorsal SUB to the MEC in rats

We next tested whether the SUB-to-MEC GABAergic projection is also present in rats by injecting AAV1-mDlx-GFP into the rat dorsal SUB (Fig. 5a, n = 4 rats). GFP-labeled SUB neurons expressed GAD67 (Fig. 5b). We observed GFP-labeled axons across all dorsal MEC layers (Figs. 5c and 5d). However, the laminar distribution unexpectedly differed from that in mice (Fig. 1); specifically, the labeled axons were most densely distributed in layer II among all other layers, whereas layers I and III also contained more labeled axons than deep layers (layers Va, Vb, and VI; Figs. 5c and 5d). GFP-labeled axonal boutons often colocalized with the vesicular GABA transporter (VGAT) in both the dorsal SUB and dorsal MEC (Fig. 5e), suggesting that these neurons form GABAergic synapses in both regions. We further assessed inhibitory synapse formation using AAV1-mDlx-SynaptoTAG2, which drives the GABAergic neuronal expression of tdTomato and EGFP-fused synaptobrevin-2 (Syb2-EGFP) to track cytoplasm and presynaptic terminals, respectively (Li et al. 2021). After injecting this AAV into the dorsal SUB (n = 2 rats), we observed presynaptic boutons triple-labeled with tdTomato, Syb2-EGFP, and VGAT in both the SUB (Fig. 5f) and MEC (Fig. 5g), further supporting that subicular GABAergic neurons form inhibitory synapses in the MEC.

**Fig. 5.**
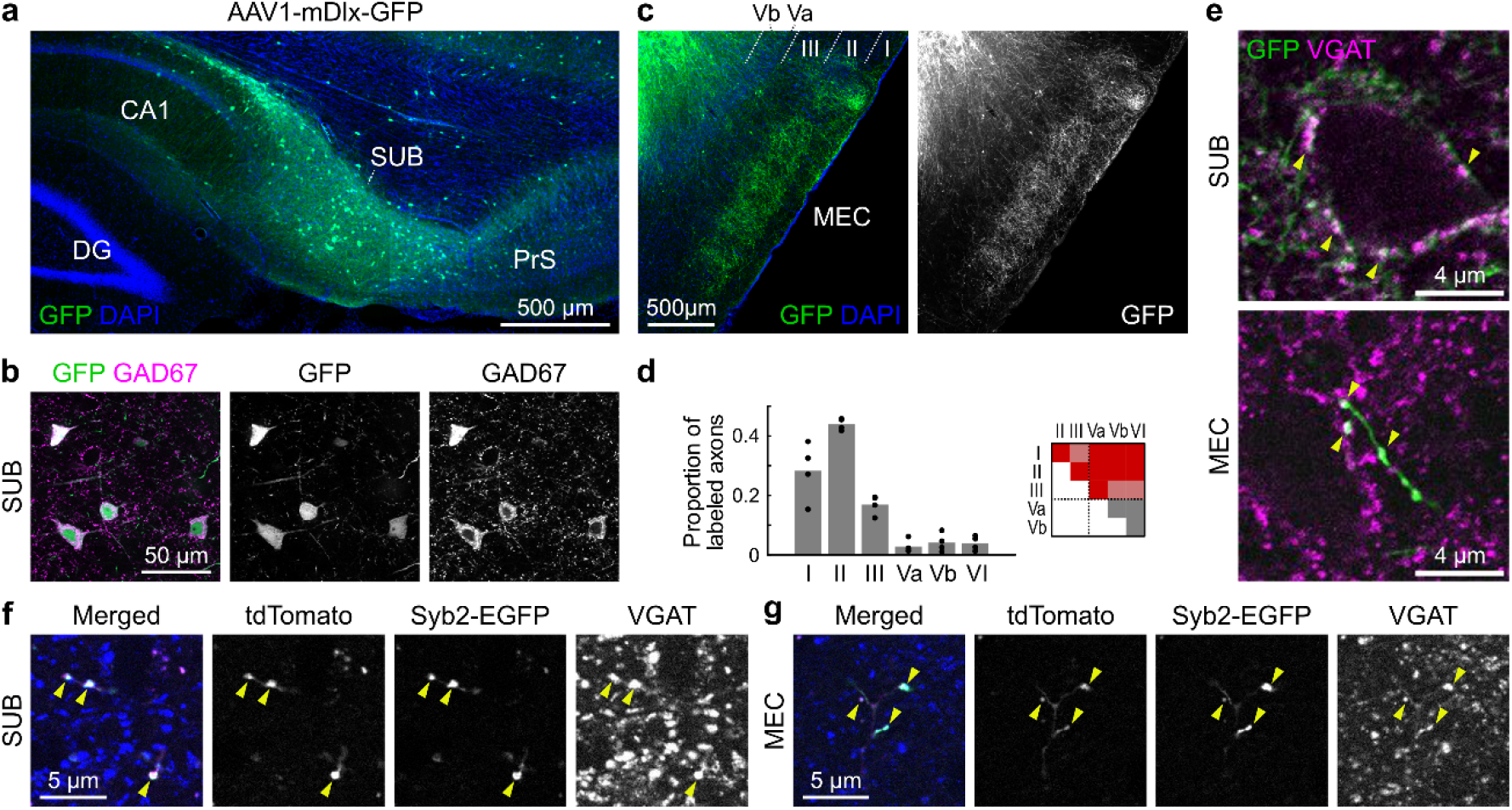
Anterograde tracing of subicular GABAergic neurons in rats. (a) Sagittal section around the rat dorsal SUB near the AAV1-mDlx-GFP injection site. (b) GFP expression in GAD67-positive SUB neurons. (c) Sagittal section of the dorsal MEC ipsilateral to the injection site, showing GFP-labeled axons. Numbers, MEC layers; dotted lines, layer boundaries. (d) Proportion of GFP-labeled axon lengths in MEC layers. Dots, individual rats; bars, means. The inset presents a *P*-value table. Red, *P* < 0.01; light red, *P* < 0.05; gray, non-significant; Tukey–Kramer test. Dotted lines, borders between superficial (I–III) and deep (Va–VI) layers. (e) Vesicular GABA transporter (VGAT) expression in GFP-labeled axons in the SUB (top) and MEC (bottom). Yellow arrowheads, colocalized expression of GFP and VGAT. (f, g) VGAT (blue) expression in tdTomato-labeled (red) and Syb2-EGFP–labeled (green) axons in the SUB (f) and MEC (g) after injection of AAV1-mDlx-SynaptoTAG2 (tdTomato-2A-Syb2-EGFP) into the dorsal SUB. Yellow arrowheads indicate boutons triple-labeled with tdTomato, Syb2-EGFP, and VGAT.

As in mice (Fig. 4), we next performed anterograde tracing of subicular PV neurons in rats (Fig. 6). After injecting AAV1-S5E2-GFP-fGFP into the rat dorsal SUB, GFP expression was observed in PV-positive neurons in the SUB (Fig. 6a). We observed GFP-labeled axons mainly in layer II of the dorsal MEC (Fig. 6b), similar to the pattern obtained by pan-GABAergic neuronal labeling (Figs. 5c and 5d). This result indicates that the dorsal SUB contains PV-positive GABAergic neurons that project to the dorsal MEC in rats, as observed in mice.

**Fig. 6.**
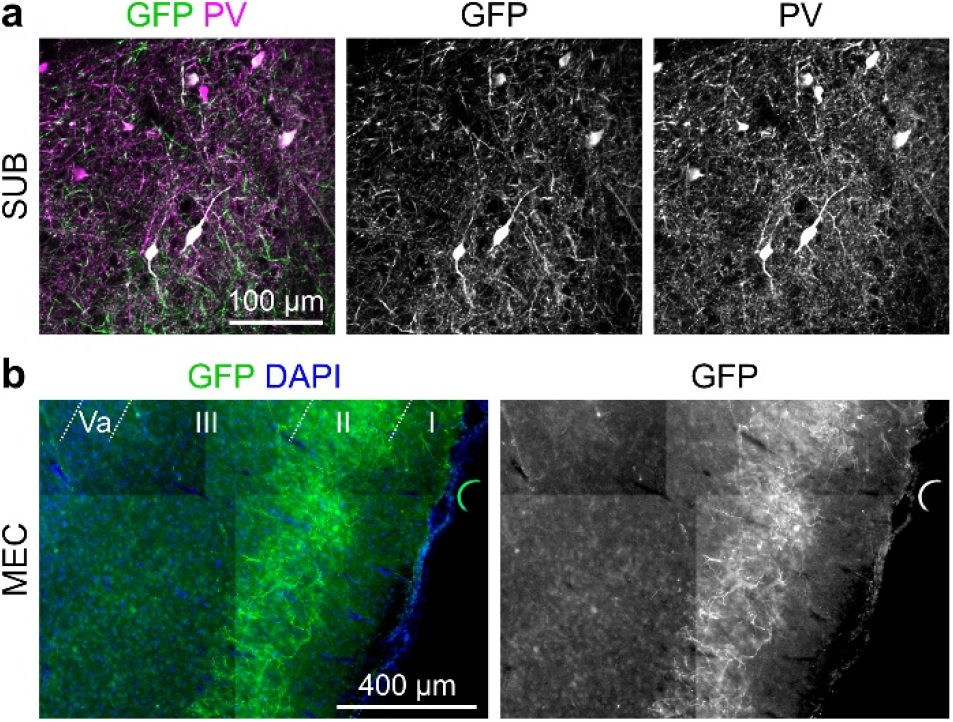
Anterograde tracing of subicular PV neurons in rats. (a) Sagittal section of the dorsal SUB near the AAV1-S5E2-GFP-fGFP injection site showing PV expression in GFP-labeled neurons. (b) GFP-labeled axons in the dorsal MEC. Numbers, MEC layers; dotted lines, layer boundaries.

## Discussion

Our data demonstrate that PV-positive GABAergic neurons in the dorsal SUB project to the dorsal MEC in both mice and rats. In contrast to subicular glutamatergic axons, which preferentially terminate in deep MEC layers, GABAergic axons were distributed across all MEC layers, with enrichment in superficial layers, and they formed inhibitory synapses. These results indicate that, in addition to excitatory neurons, inhibitory neurons also contribute to subicular output to the MEC.

Specific labeling of MEC-projecting GABAergic neurons using the dual AAV approach allowed us to identify their neuronal subtype. Immunohistochemistry indicated that a subset of these neurons expressed PV, whereas SST-positive neurons were infrequent. The remaining GABAergic projection neurons may include additional subtypes, such as VIP-, Lamp5-, or Sncg-expressing cells (Yao et al. 2021; Tzilivaki et al. 2023). A recent study identified a subset of subicular PV-positive neurons that project to the anteroventral thalamic nucleus and medial mammillary body (Baccini et al. 2025). Thus, PV-positive SUB neurons comprise multiple classes of projection neurons. Importantly, our finding that PV-positive SUB neurons project to the MEC is informative regarding potential off-target labeling. A GABAergic projection from the hippocampal CA1 area to the MEC arises from SST-positive, but not PV-positive, neurons (Melzer et al. 2012). Therefore, the potential spread of the viral solution into the CA1 area cannot explain the axonal distribution of the PV-specific AAV1-S5E2-GFP-fGFP. The GABAergic neuron subtypes of MEC-projecting presubicular neurons have not been established (van Haeften et al. 1997), leaving the possibility that multiple inhibitory sources to the MEC exist with distinct cellular compositions.

Intriguingly, the axonal distributions of subicular GABAergic neurons differed between mice and rats. In mice, the labeled axons were preferentially distributed in the superficial layers (I–III) of the MEC, whereas in rats they were enriched in MEC layer II. For excitatory projections, the overall projection patterns of the hippocampal–entorhinal circuits are similar between mice and rats, whereas species differences emerge in fine-scale connectivity (Deller et al. 2007). For instance, the entorhinal cortex projects to the contralateral dentate gyrus in rats, but not in mice (van Groen et al. 2002). To the best of our knowledge, this is the first study to describe a difference in a long-range GABAergic projection between mice and rats. Because quantitative comparisons across species are constrained by unavoidable methodological differences (e.g., injection volume, viral spread, tropism), further studies are required to validate this difference.

The functional roles of the subicular GABAergic projection to the MEC remain an open question. The SUB exhibits oscillatory activity paced at theta, gamma, and sharp-wave/ripples (Chrobak and Buzsáki 1996; Jackson et al. 2011; Jackson et al. 2014; Kitanishi *et al*. 2021; Mizuseki and Kitanishi 2022). GABAergic projections might transmit these rhythms to the MEC, thereby shaping communication between the SUB and MEC. In the MEC, superficial layers constitute a major output to the hippocampus, whereas deep layers are major targets of hippocampal/subicular excitatory input. Because the GABAergic projection described in this study preferentially innervates the MEC superficial layers, synchronous activation of SUB excitatory and inhibitory outputs could transmit SUB information via excitatory pathways while suppressing MEC-to-hippocampus/SUB signaling. These hypotheses await future investigation.

## Methods

### Ethics statement

All animal care and experimental procedures were approved by the Animal Experiment Ethics Committee of the Graduate School of Arts and Sciences at the University of Tokyo (approval no. 2022-3) and the Center for Laboratory Animal Research at Tohoku University (approval no. 2025LsA-006). The experiments were conducted in accordance with the National Institutes of Health Guide for the Care and Use of Laboratory Animals.

### Surgery for AAV-based tracing

Stereotaxic injections were performed in male C57BL/6J mice (Japan SLC, Shizuoka, Japan) at 8–9 weeks of age or male Long-Evans rats (Japan SLC) at 8–10 weeks of age. Mice were anesthetized via an intraperitoneal injection of a mixture of medetomidine hydrochloride (0.3 mg/kg), midazolam (4 mg/kg), and butorphanol tartrate (5 mg/kg) (Kitanishi and Matsuo 2017; Kitanishi et al. 2022). Rats were anesthetized with 1.5%–2% isoflurane delivered in a 1:1 (vol/vol) mixture of air and oxygen (total flow rate, 0.4 L/min) (Kitanishi et al. 2015; Kitanishi et al. 2021). The animal’s head was secured in a stereotaxic frame (Model 962, Kopf Instruments, Tujunga, CA, USA). After exposing the skull, small craniotomies were made dorsal to the injection sites using a drill (Model 1474, Kopf Instruments). AAVs were pressure-injected through a pulled glass pipette (tip diameter, 40–70 µm; 3-000-203-G/X, Drummond Scientific, Broomall, PA, USA; PC-100, Narishige, Tokyo, Japan) using a microinjector (3-000-207, Nanoject III, Drummond Scientific) at a rate of 1 nL/s. The pipette was left in place for 5 min before withdrawal. Postoperatively, animals received a single dose of meloxicam (1 mg/kg, s.c.) and cefazolin (50 mg/kg, i.m.).

Injection coordinates were as follows: mouse left dorsal SUB (AP −3.4, ML 2.1, DV 1.4), mouse left dorsal MEC (AP 0.0, ML 3.1, DV 2.0), and rat bilateral dorsal SUB (AP −5.9, ML 2.3, DV 3.0, and AP −6.1, ML 3.4, DV 3.0 for each hemisphere). AP, ML, and DV indicate the coordinates of the injection sites (in millimeters) in the anteroposterior direction from the bregma (for the SUB) or from the lambdoid suture (for the MEC), that in the mediolateral direction from the midline, and that in the dorsoventral direction from the cortical surface, respectively.

The following AAVs were used: AAV1-mDlx-GFP (83900-AAV1, Addgene, Watertown, MA, USA; ≥7.0 × 10^12^ vg/mL; 50–100 nL for mice; 250–300 nL for rats) (Dimidschstein et al. 2016), AAV1-CaMKIIα-mCherry (114469-AAV1, Addgene; ≥7.0 × 10^12^ vg/mL; mixed at 1:1 volume ratio with AAV1-mDlx-GFP; 40–60 nL in total), AAV6-pgk-Cre (24593, Addgene; 2.4 × 10^12^ vg/mL; 30–180 nL) (Kitanishi and Matsuo 2017), AAV1-hDlx-Flex-GFP (83895-AAV1, Addgene; ≥7.0 × 10^12^ vg/mL; 40–100 nL) (Dimidschstein et al. 2016), AAV1-S5E2-GFP-fGFP (135631-AAV1, Addgene; 2.5 × 10^13^ vg/mL; 50 nL) (Vormstein-Schneider et al. 2020), AAV1-Sst44-GFP (172310, Addgene; 1.8 × 10^13^ vg/mL; 400 nL) (Hrvatin et al. 2019), AAV-PHP.eB-BiSSTe4-dTomato-nlsdTomato (213944-PHPeB, Addgene; 2.3 × 10^13^ vg/mL; 50–100 nL; 1:1–1:1000) (Furlanis et al. 2025), and AAV1-mDlx-SynaptoTAG2 (7.4 × 10^12^ vg/mL, 250–300 nL).

The pAAV1-mDlx-SynaptoTAG2 plasmid (250438, Addgene) was constructed by replacing the hSyn promoter sequence in the pAAV-SynaptoTAG2 plasmid (175275, Addgene) (Li et al. 2021) with the mDlx enhancer sequence from the pAAV-mDlx-GFP plasmid (83900, Addgene) using the *Mlu*I and *Bam*HI restriction sites introduced at the ends of the mDlx sequence. The resulting construct was packaged as AAV1 as described previously (Kitanishi et al. 2022). To minimize the risk of cross-contamination, AAV6-pgk-Cre and AAV1-hDlx-Flex-GFP were injected separately on different days separated by a 6–8-day interval. We tested AAV1-Sst44-GFP and AAV-PHP.eB-BiSSTe4-dTomato-nlsdTomato for SST neuron-selective labeling (Hrvatin et al. 2019; Furlanis et al. 2025); however, these AAVs exhibited insufficient selectivity when locally injected into the SUB in our preliminary experiments.

### Histology

Animals were transcardially perfused with saline followed by 4% paraformaldehyde (PFA) in 0.1 M phosphate buffer under isoflurane anesthesia 14–16 days after the last surgery. Brains were removed and postfixed in the same fixative overnight at 4°C. They were then transferred to 30% sucrose in phosphate-buffered saline (PBS) for more than 48 h at 4°C and sectioned at a thickness of 50 µm in either the sagittal or horizontal plane using a freezing microtome (SM2010R, Leica, Nussloch, Germany; EF-13, Nihon Microtome Laboratory, Osaka, Japan). Free-floating sections were incubated sequentially in 5% bovine serum albumin (BSA)/0.3% Triton X-100 in PBS for 30 min at room temperature, primary antibodies diluted in 5% BSA/PBS overnight at either 4°C or room temperature, and corresponding secondary antibodies together with either DAPI (0.5 µg/mL, D1306, Thermo Fisher Scientific, Waltham, MA, USA) or NeuroTrace 530/615 Red Nissl Stain (N21482, Thermo Fisher Scientific) in 5% BSA/PBS for either 30 min or 2 h at room temperature. Sections were washed with PBS between incubations. The following primary antibodies were used: chicken anti-GFP (1:1000, ab13970, Abcam, Cambridge, UK), mouse anti-GAD67 (1:1000, MAB5406, Sigma-Aldrich, St. Louis, MO, USA), rabbit anti-PV (1:1000, PV27a, Swant, Fribourg, Switzerland), rabbit anti-SST (1:1000, T4103, BMA Biomedicals, Augst, Switzerland), and rabbit anti-VGAT (1:1000, MSFR106110, Nittobo Medical, Tokyo, Japan). The secondary antibodies (all from Thermo Fisher Scientific) were as follows: goat anti-chicken IgY conjugated with Alexa Fluor 488 (1:200, A-11039), goat anti-mouse IgG conjugated with Alexa Fluor 594 (1:800, A-11032), goat anti-rabbit IgG conjugated with Alexa Fluor 594 (1:800, A-11037), and goat anti-rabbit IgG conjugated with Alexa Fluor 647 (1:500, A-21245). Sections were mounted on glass slides with antifade mounting medium (P36961 or 00-4958-02, Thermo Fisher Scientific). Images were acquired using fluorescence microscopes (Axio Imager 2 with Apotome 3, Zeiss, Oberkochen, Germany; A1R, Nikon, Tokyo, Japan).

The lengths of GFP-labeled axons in each MEC layer were quantified by manual tracing of axon segments in sagittal section images using Fiji software (ver. 1.54p). Total axon length was summed for each layer, and the proportion of labeled axons was calculated.

Cells positive for GFP, GAD67, PV, and SST were counted, and the superficial and deep boundaries of the SUB cell layer were outlined using Fiji software. Normalized positions within the SUB cell layer were determined using a custom-written MATLAB script (R2025b, MathWorks, Natick, MA, USA). The superficial and deep boundary curves were sampled at 200 evenly spaced points along the proximal–distal axis. Corresponding points were connected to generate transverse lines, which were then subdivided into 200 equal segments along the superficial–deep axis, yielding a 200 × 200 grid across the cell layer. A cell’s normalized position was defined by the grid indices of the bin containing that cell.

Statistical analyses were performed using MATLAB. All statistical tests were two-sided, and statistical significance was assessed at an alpha level of 0.05.

### Surgery for ex vivo electrophysiology

Male C57BL/6J mice (Japan SLC) at 6–7 weeks of age were anesthetized with isoflurane in an induction chamber and subsequently injected intraperitoneally with ketamine (80 mg/kg) and xylazine (10 mg/kg). Animals were mounted on a stereotaxic frame, and isoflurane was continuously delivered via a surgical anesthesia mask at 1%–2% throughout the procedure. Eye ointment was applied to prevent corneal drying. The skin overlying the skull was disinfected with iodide, and local anesthesia was administered subcutaneously (lidocaine, 10 mg/kg). After exposing the skull, a small burr hole was drilled above the injection site. AAV9-mDlx-ChR2-mCherry (1.2 × 10^13^ GC/mL; 60 nL; 83898, Addgene) was injected into the dorsal SUB at a rate of 12 nL/min using a 1-µL Hamilton microsyringe. The injection pipette was left in place for an additional 10 min before withdrawal to prevent backflow. The incision was sutured, and the animals were returned to their home cages after recovery from anesthesia.

### Acute slice preparation

Two to three weeks after viral injection, mice were deeply anesthetized with isoflurane and decapitated. Brains were rapidly removed and immersed in ice-cold oxygenated cutting solution containing the following (in mM): 110 choline-Cl, 2.5 KCl, 7 MgCl_2_, 0.5 CaCl_2_, 25 glucose, 1.25 NaH_2_PO_4_, 25 NaHCO_3_, 11.5 Na-ascorbate, 3 Na-pyruvate, and 100 D-mannitol. Sagittal slices (350 µm thick) containing MEC were prepared using a vibratome (DSK LinearSlicer PRO 7, DSK, Kyoto, Japan) while maintaining the tissue submerged in ice-cold cutting solution. Slices were transferred to a holding chamber and incubated at 35°C in an oxygenated holding solution containing the following (in mM): 126 NaCl, 3 KCl, 1.2 NaH_2_PO_4_, 10 glucose, 26 NaHCO_3_, 3 MgCl_2_, and 0.5 CaCl_2_. After 30 min of incubation, slices were maintained at room temperature for at least 30 min before recording.

### Electrophysiological recording

Acute brain slices were transferred to a recording setup and visualized using infrared differential interference contrast optics with a ×20/1.0 NA water immersion objective (Axio Examiner D1, Zeiss). Recordings were performed at 28°C–30°C in oxygenated artificial cerebrospinal fluid (ACSF) containing the following (in mM): 126 NaCl, 3 KCl, 1.2 NaH_2_PO_4_, 10 glucose, 26 NaHCO_3_, 1.5 MgCl_2_, and 1.6 CaCl_2_. Whole-cell voltage–clamp recordings were obtained using borosilicate glass pipettes (2–4 MΩ) filled with a Cs-based internal solution containing the following (in mM): 120 Cs-gluconate, 10 HEPES, 10 Na-phosphocreatine, 4 Mg-ATP, 0.4 Na-GTP, 0.5 EGTA, 2 QX314, and 10 TEA-chloride. The internal solution was adjusted to pH 7.3 and 290 mOsm. Biocytin (2.5%) was included in the internal solution, which was allowed to diffuse throughout the cell for post hoc identification of the soma location and neuronal morphology.

Electrophysiological signals were acquired using an Integrated Patch Amplifier with dual headstage (Double IPA, Sutter Instrument, Novato, CA, USA) and SutterPatch software (Sutter Instrument), low-pass filtered at 2 kHz, and digitized at 10 kHz. Holding potentials in the voltage– clamp mode were corrected for a liquid junction potential of approximately −15 mV. Accordingly, excitatory and inhibitory postsynaptic currents (EPSCs and IPSCs) were recorded at holding potentials of −50 and +20 mV, respectively, corresponding to the reversal potentials for GABA-A receptors (−65 mV) and AMPA receptors (+5 mV). In some experiments, monosynaptic inputs were isolated by adding tetrodotoxin (TTX, 1 µM), 4-aminopyridine (4-AP, 100 µM), and elevated Ca^2+^ (4 mM) to the ACSF, which blocks action potential propagation while permitting neurotransmitter release from directly photoactivated presynaptic terminals. To pharmacologically confirm GABAergic transmission, the competitive GABA-A receptor antagonist gabazine (10 µM) was applied. All chemicals were obtained from Sigma-Aldrich or FUJIFILM Wako Pure Chemical Industries (Osaka, Japan). Data were excluded if the series resistance exceeded 40 MΩ.

Channelrhodopsin-2 (ChR2)-expressing presynaptic axon terminals in the MEC were activated by brief blue light pulses (475 nm) delivered from an LED light source (Colibri 7, Zeiss) through the ×20/1.0 NA objective (Axio Examiner D1). The light pulse power and duration were typically set to 38.6 mW and 2 ms, respectively.

EPSC and IPSC events were analyzed offline using the Synaptic Event Analysis module from the SutterPatch application (Igor Pro, WaveMetrics, Portland, OR, USA). Only slices with at least two successful synaptic responses to photostimulation were included in the analysis. No outliers were removed from the data.

All data presented in the figures are shown as mean ± standard error of the mean. R (version 2026.01.0, The R Foundation for Statistical Computing, Vienna, Austria) was used for data analysis, and one-way ANOVA with Bonferroni’s multiple comparison test was used to compare the amplitudes of IPSCs between cell types. Thresholds for significance were set at \**P* < 0.05, \*\**P* < 0.01, and \*\*\**P* < 0.001.

### Histology and imaging of the recorded slices

After electrophysiological recordings, brain slices were fixed overnight at 4°C in 4% PFA. Slices were washed five times for 15 min each in PBS containing 0.3% Triton X-100 (PBS-Tx), followed by incubation in a blocking solution containing PBS-Tx supplemented with 10% normal goat serum for 3 h at room temperature. To visualize subicular GABAergic neurons expressing ChR2-mCherry and identify the cytoarchitecture, slices were incubated with primary antibodies against DsRed (rabbit anti-DsRed, 1:500, 632496, Takara Bio, Tokyo, Japan) and NeuN (mouse anti-NeuN, 1:500, MAB377, Chemicon), diluted in blocking solution for 4 days at 4°C. Slices were washed five times for 15 min each in PBS-Tx and incubated overnight at room temperature with secondary antibodies diluted in PBS-Tx, namely goat anti-rabbit Cy3 (1:200, 111-165-144, Jackson ImmunoResearch, West Grove, PA, USA) and goat anti-mouse Alexa Fluor® 488 (1:200, 115-545-146, Jackson ImmunoResearch). Fluorescently conjugated streptavidin (Alexa Fluor® 647 Streptavidin, 1:200, 016-600-084, Jackson ImmunoResearch) was included during secondary antibody incubation to visualize biocytin-filled recorded neurons. After incubation, the slices were washed three times for 15 min each in PBS-Tx and stored in PBS at 4°C until imaging.

For imaging, slices were mounted in custom-made metal well slides, coverslipped, and imaged using a laser-scanning confocal microscope (LSM 900, Zeiss). Low-magnification overview images were acquired using a Plan Apochromat ×10 objective (NA 0.45) to confirm the laminar location of recorded neurons, and higher-magnification images were obtained using a Plan Apochromat ×20 objective (NA 0.8) to assess neuronal morphology. Both overview and high-magnification images were acquired encompassing the full extent of each recorded neuron to reconstruct complete cellular morphology.

## Author contributions

Conceptualization: T.A. and T.K.; Investigation: all authors; Supervision: T.K; Writing, original draft: S.O. and T.K.; Writing, review, and editing: all authors; Funding acquisition: T.A., S.O., and T.K.

## Data availability

The data are available from the corresponding author upon request. Source data are provided with this paper.

